# Synchronization and interaction of proline, ascorbate and oxidative stress pathways under abiotic stress combination in tomato plants

**DOI:** 10.1101/2020.06.30.179770

**Authors:** María Lopez-Delacalle, Christian J Silva, Teresa C Mestre, Vicente Martinez, Barbara Blanco-Ulate, Rosa M Rivero

## Abstract

Adverse environmental conditions have a devastating impact on plant productivity. In nature, multiple abiotic stresses occur simultaneously, and plants have evolved unique responses to cope against this combination of stresses. Here, we coupled genome-wide transcriptional profiling and untargeted metabolomics with physiological and biochemical analyses to characterize the effect of salinity and heat applied in combination on the metabolism of tomato plants. Our results demonstrate that this combination of stresses causes a unique reprogramming of metabolic pathways, including changes in the expression of 1,388 genes and the accumulation of 568 molecular features. Pathway enrichment analysis of transcript and metabolite data indicated that the proline and ascorbate pathways act synchronously to maintain cellular redox homeostasis, which was supported by measurements of enzymatic activity and oxidative stress markers. We also identified key transcription factors from the basic Leucine Zipper Domain (bZIP), Zinc Finger Cysteine-2/Histidine-2 (C2H2) and Trihelix families that are likely regulators of the identified up-regulated genes under salinity+heat combination. Our results expand the current understanding of how plants acclimate to environmental stresses in combination and unveils the synergy between key cellular metabolic pathways for effective ROS detoxification. Our study opens the door to elucidating the different signaling mechanisms for stress tolerance.

**HIGHLIGHTS:**

- The combination of salinity and heat causes a unique reprogramming of tomato metabolic pathways by changing the expression of specific genes and metabolic features.
- Proline and ascorbate pathways act synchronously to maintain cellular redox homeostasis
- Key transcription factors from the basic Leucine Zipper Domain (bZIP), Zinc Finger Cysteine-2/Histidine-2 (C2H2) and Trihelix families were identified as putative regulators of the identified up-regulated genes under salinity and heat combination.

## INTRODUCTION

Multiple environmental factors such as salinity, high temperatures, cold, or drought cause abiotic stresses in plants, which result in large agricultural losses worldwide, estimated to be around $14–19 million (Rivera *et al*., 2017). Under field conditions, different abiotic stressors usually occur at the same time; for example, it is common that high temperatures coexist with highly saline soils or water scarcity. Studies in the last decade have shown that plant responses to combined abiotic stresses are unique and cannot be deduced from the study of plants subjected to each stress separately (Mittler, 2006; Miller *et al*., 2010; Rivero *et al*., 2014; Anjum *et al*., 2019; Sehgal *et al*., 2019; Lopez-Delacalle *et al*., 2020; Zandalinas *et al*., 2020).

Many metabolic mechanisms act in concert during the plant’s response to abiotic stress, including rapid changes in gene expression, ionic adjustment, and activation and inactivation of proteins that carry out the synthesis and degradation of compounds used for cell signaling and protection (e.g., osmoprotectants and antioxidants), among others (Rivero *et al*., 2014; Zushi *et al*., 2014; Zandalinas *et al*., 2020). Proline has been widely reported to act as an osmoprotectant in plant defense against certain stress conditions, such as drought and salinity (Rivero *et al*., 2004*b*, 2014)(Rivero *et al*., 2004*b*, 2014; Martinez *et al*., 2018). Under heat stress, plants synthesize proline, as reported by the induction of pyrroline-5-carboxylate synthase (P5CS) and the subsequent accumulation of the amino acid (Rivero *et al*., 2004*b*; Torres *et al*., 2006). Shalata & Neumann (2001) have also reported that under salinity, proline accumulation can improve plant salt tolerance and reduce oxidative damage by decreasing lipid peroxidation in tomato plants.

Stress conditions cause the accumulation of reactive oxygen species (ROS), which are known to induce oxidative stress (e.g., lipid peroxidation) and serve as signaling molecules in plants (Suzuki *et al*., 2012; Kollist *et al*., 2019). Plants accumulate antioxidants, such as ascorbate (ASC), glutathione (GSH), carotenoids, tocopherols, and flavonoids, and activate enzymatic reactions to maintain cell homeostasis under increasing oxidative conditions. The ASC/GSH cycle is critical for detoxifying ROS from plant cells. Briefly, the hydrogen peroxide (H_2_O_2_) generated by detoxification of superoxide radicals is further converted to H_2_O and O_2_ by ASC peroxidase (APX), the first enzyme of the ASC/GSH cycle, using ASC as an electron donor (Noctor and Foyer, 1998). Because ASC is considered the first antioxidant line of defense in H_2_O_2_ detoxification (Foyer and Noctor, 2011; Akram *et al*., 2017), it is expected that plants with a high cellular accumulation of this compound will have a greater tolerance to oxidative stress. In fact, tomato seeds treated with ascorbic acid have been shown to have better tolerance to salinity stress, improved germination, and better growth parameters (Sayed, 2013). We have previously reported (Rivero et al. 2004) that enzymes that belong to the ASC/GSH cycle were inhibited in tomato plants under high temperature, leading to H_2_O_2_ accumulation and inhibition of plant growth and yield. In addition to its importance in ROS detoxification, the cellular content of GSH and ASC improves osmoregulation, efficient use of water, photosynthetic activity, and general parameters of plant productivity (Noctor and Foyer, 1998; Meyer, 2008; Foyer and Noctor, 2011).

Plants need to rapidly regulate and fine-tune their responses to stress in order to maximize energy expenditure in adverse conditions. Transcription factors (TFs) are considered key components in the control of abiotic stress signaling (Schmidt *et al*., 2012; Castelán-Muñoz *et al*., 2019); however, little is known about their role in stress combination. Just recently, a report by Zandalinas et al. (Zandalinas *et al*., 2020) found that Arabidopsis plants induced a unique set of TFs when subjected to different abiotic stress combinations, and that those genes were relatively unique across stress conditions.

Here we hypothesize that the combination of salinity and heat induces a unique physiological response in tomato plants by activating specific regulatory and metabolic pathways that act synergistically to maintain cell redox homeostasis. In this work, we analyze how the combination of salinity and heat affects the transcriptome and metabolome of tomato plants in order to find the unique elements that are differentially regulated under these stress conditions and that may be key in ROS detoxification and, thus, plant tolerance to abiotic stress combination.

## MATERIALS AND METHODS

### Plant material, experimental design, and growth conditions

*Solanum lycopersicum* L cv. Boludo (Monsanto) seeds were germinated in vermiculite under optimal and controlled conditions in a growth chamber (chamber A). These conditions were: a photoperiod of 16/8 hours of day/night with a light intensity of 500 µmol m^−2^ s^−1^, a relative humidity (RH) between 60 and 65% and a temperature of 25 °C. Subsequently, when the plants had at least two true leaves, they were transplanted to an aerated hydroponic system containing a modified Hoagland solution and grown under these conditions for one week. The nutrient solution had the following composition: KNO_3_ (3 mM), Ca(NO_3_)_2_ (2 mM), MgSO_4_ (0.5 mM), KH_2_PO_4_ (0.5 mM), Fe-EDTA (10 µM), H_3_BO_3_ (10 µM), MnSO_4_·H_2_O (1 µM), ZnSO_4_·7H_2_O (2 µM), CuSO_4_·5H_2_O (0.5 µM), and (NH_4_)_6_Mo_7_O_24_·4H_2_O (0.5 µM) (Hoagland and Arnon, 1950). The electric conductivity (EC) and pH of the nutrient solution were measured and maintained within 1.4–1.7 mS-cm^-1^ and 5.2–5.6, respectively. After acclimatization, half of the plants were transferred to a twin chamber whose temperature was previously set at 35 °C (chamber B). In both twin chambers, a saline concentration in the nutrient solution of 75 mM NaCl was added to half of the plants. Therefore, four different conditions were obtained: control (25 °C 0 mM NaCl), salinity (25 °C and 75 mM NaCl), heat (35 °C 0 mM NaCl), and salinity and heat (35 °C and 75 mM NaCl). Plants were kept under these conditions for a period of 14 days. After this time, six plants from each treatment were sampled for subsequent analysis.

### Measurements of photosynthetic parameters

Photosynthetic parameters were determined on a fully-expanded, metabolically-mature middle leaf in all plants. These data were taken with a gas exchange system (LI-COR 6400, Li-Cor) at the beginning (day 0), the middle (day 7), and at the end of the experiment (day 14). The conditions established in the LI-COR were: 1000 µmol photons m^-2^ s^-1^ and 400 µmol mol^-1^ CO_2_. The leaf temperature was maintained at 25 °C for control and salinity treatment plants, and at 35 °C for plants in the high temperature, and the combination of high temperature and salinity treatments. The leaf-air vapor pressure deficit was maintained between 1-1.3 kPa. With this analysis, the device reported data on CO_2_ assimilation, stomatal conductance, and transpiration rate. At the end of the experiment, the leaves, stem, and root were separated, and the fresh weight (FW) of each part of the plant was determined separately.

### Quantification of oxidative stress-related markers

#### H_2_O_2_ accumulation

H_2_O_2_ was extracted as described by Yang et al. (2007), with some modifications, which are fully described in García-Martí et al. (García-Martí *et al*., 2019). The samples were used for the future determination of H_2_O_2_ concentration and lipid peroxidation. H_2_O_2_ was quantified as described by MacNevin and Urone (1953).

#### Lipid peroxidation

For lipid peroxidation determination, malondialdehyde (MDA) was used, which is a product of the peroxidation of membrane lipids. The same enzyme extract as the one utilized for the determination of H_2_O_2_ was used. The procedure was described by Fu and Huang (2001).

#### Antioxidant capacity

Regarding antioxidant capacity, it was carried out according to the protocol by Koleva *et al*. (2002). The remaining amount of 2,2-diphenyl-1-picrylhydrazyl (DPPH), measured at a certain time, is inversely proportional to the antioxidant capacity of the substances present in the sample. Results are expressed as % Radical Scavenging Activity (RSA), or percentage of free radical scavenging activity.

#### Protein oxidation

Protein oxidation was assayed as according to Reznick and Packer (1994). PCO groups react with 2,4-dinitrophenylhydrazine (DNPH) to generate chromophoric dinitrophenylhydrazones, which can be recorded with a spectrophotometer. The absorbance was measured at 360 nm, using the molar extinction coefficient of DNPH 2.2× 10^4^ M^-1^ cm^-1^.

### RNA extraction and sequencing

Total RNA was extracted from 1 g of frozen tomato leaves using TRI-Reagent (Sigma-Aldrich) and following the manufacturer’s instruction. The quantity and quality of RNA were determined using a NanoDrop 3300 fluorospectrometer (Thermo Scientific Instruments, USA). Three biological replications for each treatment were used for total RNA extraction and sequencing. For each RNA sample, mRNA was enriched using a Dynabeads mRNA purification kit (Invitrogen), then the samples were sent to BGI-Shenzhen (hereafter ‘BGI’, China) for RNA sequencing. Sequencing was carried out on a HiSeq2000 according to the Illumina protocols for 90 □ × □ 2 pair-end sequencing covering a read length of 100 bp. An average of 10 Gb clean data per sample was generated after filtering to ensure a complete set of expressed transcripts with sufficient coverage and depth for each sample.

### Bioinformatics pipeline

#### RNA sequencing and data processing

Raw sequencing reads were trimmed for quality and adapter sequences using Trimmomatic v0.33 (Bolger *et al*., 2014) were used with the following parameters: maximum seed mismatches = 2, palindrome clip threshold = 30, simple clip threshold = 10, minimum leading quality = 3, minimum trailing quality = 3, window size = 4, required quality = 15, and minimum length = 36. Trimmed reads were mapped using Bowtie2 (Langmead and Salzberg, 2012) to the tomato transcriptome (SL4.0 release; http://solgenomics.net). Count matrices were made from the Bowtie2 results using sam2counts.py v0.91 (https://github.com/vsbuffalo/sam2counts/). A summary of the quality assessment and mapping results can be found in **Supplemental Table S1**. The raw sequencing reads and the read mapping count matrices are available in the National Center for Biotechnology Information Gene Expression Omnibus database under the accession GSE152620.

#### Differential expression analysis

The Bioconductor package DESeq2 (Love *et al*., 2014) was used to perform normalization of read counts and differential expression analyses for various treatment comparisons. Differentially expressed (DE) genes for each comparison were those with an adjusted p-value of less than or equal to 0.05.

#### Functional annotation and enrichment analyses

Basic functional annotations for genes were determined with the Automated Assignment of Human Readable Descriptions (AHRD) provided in the SL4.0 build of the tomato genome. KEGG annotations were determined using the KEGG Automatic Annotation Server (Moriya *et al*., 2007). Enrichments were conducted via Fisher’s exact test with p-values adjusted using the Benjamini and Hochberg method (Benjamini and Hochberg, 1995).

#### Promoter motif analysis

Binding motifs for tomato transcription factors were obtained from the Plant Transcription Factor Database (http://planttfdb.cbi.pku.edu.cn/). Promoter sequences defined as 1000 base pairs upstream from the transcription start site of each gene were obtained using the ‘flank’ function in bedtools v2.29.2 (https://bedtools.readthedocs.io/en/latest/). Enrichment of transcription factor binding motifs on the promoter sequences of up-regulated genes was performed using the Analysis of Motif Enrichment tool in MEME-Suite (McLeay and Bailey, 2010) using all non-up-regulated tomato genes as the control sequences, the average odds score scoring method and Fisher’s exact test.

### Metabolomics analysis

Six biological replications of frozen tomato leaves per treatment were used for the metabolomics analysis. One gram of this frozen plant material was extracted in methanol: water (3:1 v/v) as described previously in Martinez *et al*. (2016). Agilent MassHunter Qualitative analysis software v 6.00 (Agilent Technologies, Palo Alto, USA) was used to obtain an initial peak processing (**Supplemental Fig. S1**). Then, XCMS online software (www.xcmsonline.scripts.edu) which incorporates CAMERA (a Bioconductor package to extract spectra, annotate isotopes and adduct peaks, among other functions) in its analysis, was implemented in our curated raw data (**Supplemental Table S2**). A second level of statistical analysis was carried out, consisting of data normalization of the peaks obtained for each treatment against the control, and a t-test followed by an ANOVA analysis. Then, log_2_ of the fold-change was calculated and all the molecular features with a P_*adj*_ adjusted greater than 0.05 and a log_2_ fold change (FC) greater than -1 or smaller than 1 were eliminated from the analyses (**Supplemental Table S3**). All the molecular features that remained after these restricted statistical analyses were compared among the different treatments applied (supplemental Table S3; Euler diagram Fig 3B).

The metabolite identification of the molecular features of interest for this study was performed using a mathematical search based on the predicted elemental composition through some of the most important open-source databases (MOTO, KNApSAcK, KOMOCS, MassBank, ARMeC and METLIN) within a mass tolerance of 10 ppm. Then, the isotope ratio (IR) and retention time (rt) from the different metabolites identified unequivocally were checked again across the different databases mentioned above. Identified metabolites that remained after this filtering were labeled accordingly and highlighted in yellow in **Supplemental Table S4**. The concentration of the compounds that showed significant differences (P_*adj*_ <0.05 and log_2_ FC >2) under salinity and heat combination as compared to control plants and which were of interest in this study were plotted in a box-and-whisker type plot using XCMS online (**Supplemental Figs. S2 and S3**).

### Enzymatic activities

#### Crude extract

All enzymatic activities were extracted according to the procedure described by Torres *et al*. (2006).

#### Superoxide dismutase (SOD)

SOD activity was assayed as described previously by McCord JM (1969). SOD activity was expressed as units of SOD (mg prot)^-1^ (min)^-1^, a unit which indicates the amount of enzyme needed to neutralize one unit of xanthine oxidase.

#### Catalase (CAT)

CAT activity was calculated using the extinction coefficient of 39.4 mM^-1^cm^-1^ as described by Aebi (1984). CAT activity was expressed as µmol of reduced H_2_O_2_ (mg prot)^-1^ (min)^-1^.

#### Ascorbate peroxidase (APX) Dehidroascorbate reductase (DHAR) and Monodehidroascorbate reductase (MDHAR)

APX, DHAR and MDHAR activities were assayed as described by Miyake and Asada (1992). The rate of reaction was calculated using a molar extinction coefficient of 2.8 mM^-1^ cm^-1^. APX activity was expressed as µmol of reduced ascorbic acid (mg prot)^-1^ (min)^-1^.

#### Glutathione peroxidase (GPX)

GPX activity was carried out using the Glutathione Peroxidase Assay Kit (Abcam, Ref. ab102530, Cambridge, UK) considering the decrease of NADPH at 340 nm, using an extinction coefficient of 6.22 mM^−1^ cm^−1^.

#### Glutathione reductase (GR)

GR activity was assayed through the non-enzymatic NADPH oxidation (Halliwell and Foyer, 1976). The activity was determined by measuring the decrease in the reaction rate at 340nm and was calculated from the 6.22 mM^−1^ extinction coefficient.

#### Protein concentration in the enzyme extract

Proteins were quantified with the Bradford method (Bradford, 1976), in which a volume of Bradford Reagent reagent (BioRAd, Catalog No. 30214) was added to an aliquot of the enzyme extract. The absolute values as well as the calculated log_2_ of the data normalized against control plants of all the enzymatic activities assayed can be found in **Supplementary Table S5**.

### Statistical analysis

Statistical analysis for FW, photosynthetic parameters, H_2_O_2_ concentration, MDA content, protein oxidation, and enzymatic activities were performed with an analysis of variance with p-value < 0.05 set as the cut-off value, as indicative of significant differences, followed by a Duncan test and a t-test when necessary. Transcriptomics and metabolomics statistical analysis was performed as described above.

## RESULTS

### Tomato plants grown under the combination of salinity showed a better performance in the photosynthetic parameters as compared to salinity alone

Eighty-four tomato plants were grown in two independent chambers using four different environmental conditions: 25 °C and 0 mM NaCl (control), 25 °C and 75 mM NaCl (salinity), 35 °C and 0 mM NaCl (heat), and 35 °C and 75 mM NaCl (salinity + heat) for 14 days (**Fig. 1A**). Fresh weight was recorded at the end of the experiment (**Fig. 1B**). Salinity and the salinity + heat resulted in a significant reduction of biomass when compared to control plants, whereas the heat-treated plants did not differ significantly from the controls. Interestingly, when salinity and heat were applied simultaneously, the growth was significantly improved with respect to salinity up to about 18%. Photosynthetic parameters were also measured at 0 days, and after 7 and 14 days after the start of the treatments, as stress physiological markers (**Figs. 1C-F**). In our experiments, CO_2_ assimilation rate was highly inhibited after 7 days under salinity as compared to control plants, with an inhibition of 50% at 7 days, and 70% at 14 days with respect to control plants (**Fig. 1C**). The other stress treatments applied (heat and salinity + heat) did not differ significantly with respect to the values obtained in control plants during the entire experiment, contrary to that observed under salinity. Control plants had a transpiration rate and a stomatal conductance that were practically constant during the entire experiment, whereas plants subjected to heat and salinity + heat treatments showed a significant increase in transpiration rate (41%) and stomatal conductance (29%) at 7 days, which was maintained until the end of the experiment (**Figs. 1D and 1E**). Contrarily, the salinity treatment led to a significant reduction in the transpiration rate and the stomatal conductance at day 7 from the start of the treatment until the end. In this regard, the salinity + heat treatment showed a significant improvement in the photosynthetic parameters as compared to salinity alone. Curiously, no differences were found between salinity, heat, and the combination of both stresses for water use efficiency (WUE, **Fig. 1F**), but all the treatments showed a significant decrease in this parameter as compared to control plants.

**Figure 1.**
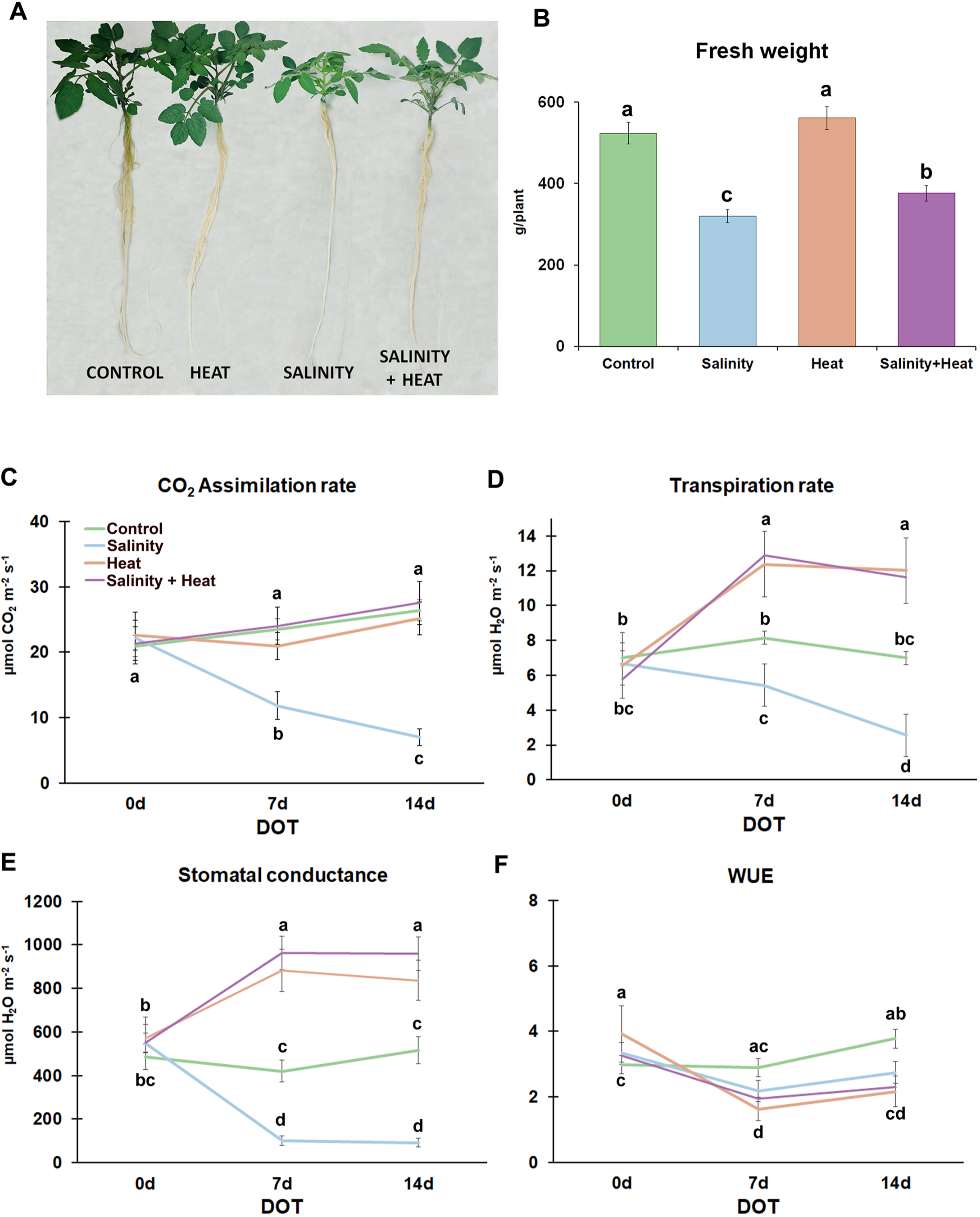
**(A)** Pictures of tomato plants at the end of the control or stress treatments. **(B)** Whole plant fresh weight (FW) of tomato plants grown under control, heat, salinity or the combination of salinity and heat. **(C-F)** Photosynthetic parameters measured in the third and four fully mature expanded leaves of tomato plants grown under control or stress conditions measured at the beginning (0 days), during (7 days) or at the end (14 days) of the treatments. Values represent means ± SE (n = 9). Bars with different letters within each panel are significantly different at p < 0.05 according to Tukey’s test.

### Salinity and heat combination induced a specific transcriptional response and pathway activation

An RNAseq study was performed to identify specific biochemical pathways or molecular functions that could explain the different physiological responses of the tomato plants to the salinity, heat, and salinity + heat treatments. RNA was sequenced from three biological replicates from each treatment, including control plants. A principal component analysis of the normalized reads revealed that all samples clustered according to treatment, which validated the unique transcriptional reprograming caused by each stress condition (**Fig. 2A**). Then, a differential expression analysis was performed to determine the individual genes affected by each treatment when compared to the control. A total of 15,852 genes were found to be differentially expressed (*P*_*adj*_ < 0.05) across all three treatments (**Supplemental Table S6**). A comparison of both up- and down-regulated genes from each of the three treatments further confirmed that each treatment resulted in a high number of unique differentially expressed genes (**Fig. 2B**). Most notably, it was found that 1,388 (7.32% of the total) were differentially expressed only for salinity + heat, with 923 genes up-regulated and 465 genes down-regulated by this stress combination (**Fig. 2B**).

**Figure 2.**
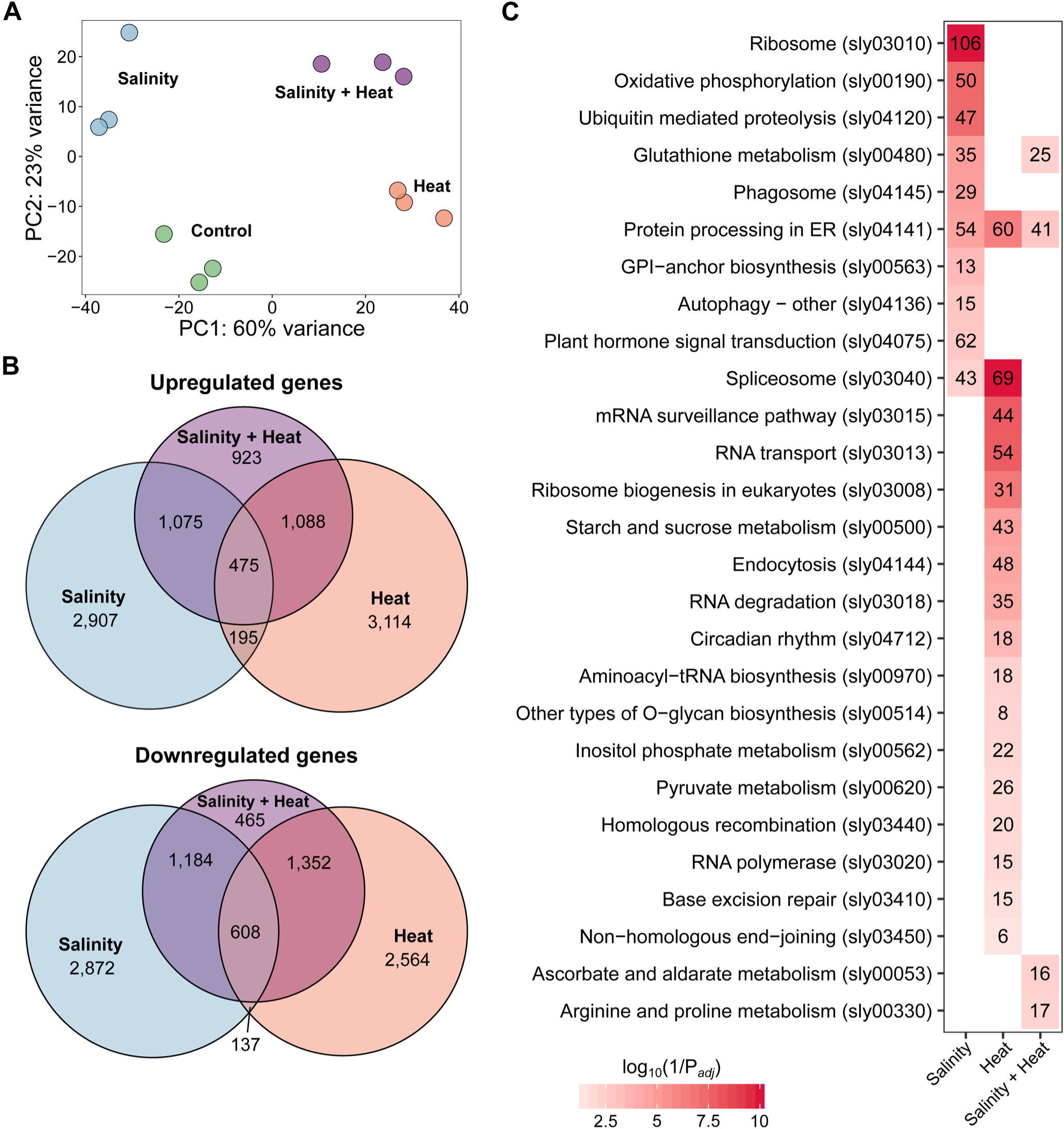
RNAseq analysis performed in tomato leaves after 14 days of growing under control or stress (salinity, heat or the combination of salinity and heat) conditions. **(A)** Principal component analysis (PCA) of the normalized reads obtained for each treatment. **(B)** Euler diagram representing up- and down-regulated genes (adjusted P<0.05) of tomato plants grown under control, simple (salinity or heat) or combined (salinity + heat) stress. **(C)** Pathway enrichment analysis performed within the up-regulated genes under the different stress conditions applied. More details on these analyses can be found in the Materials and Method section.

**Figure 3.**
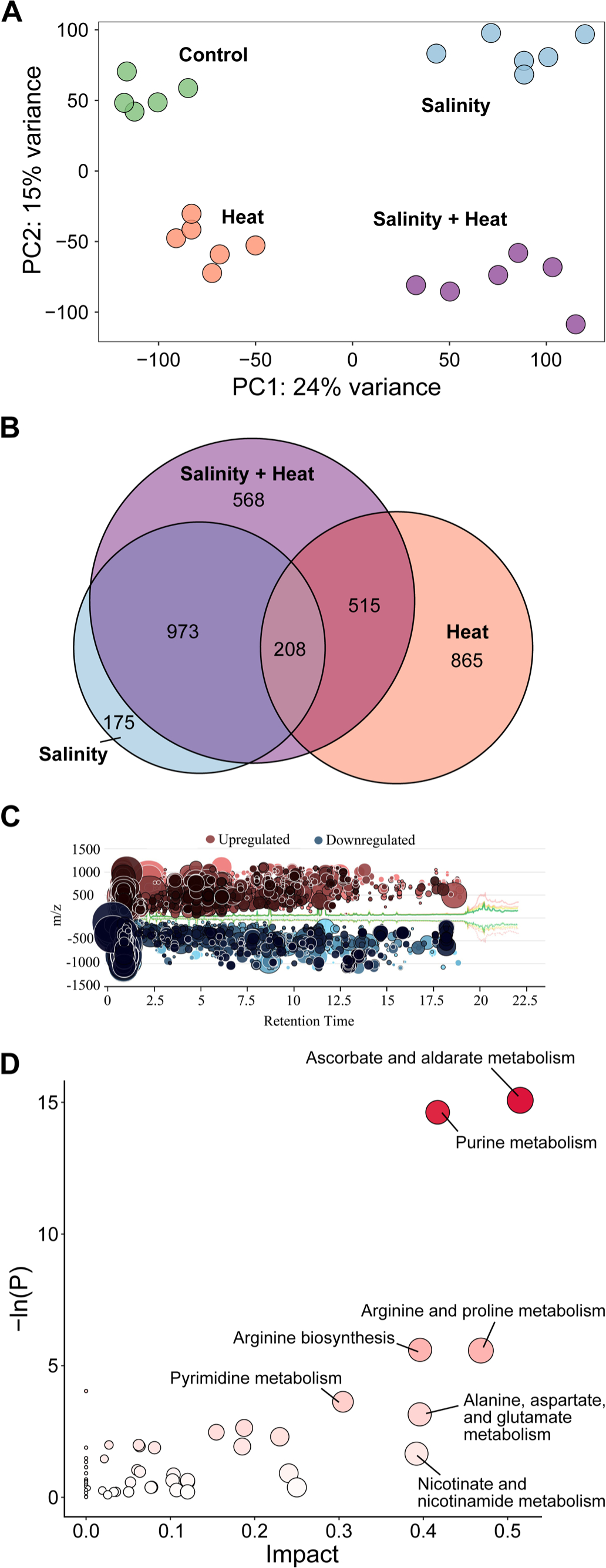
Metabolomic analysis performed in tomato leaves after 14 days of growing under control, simple (salinity or heat) or combined (salinity + heat) stress conditions. **(A)** Principal component analysis (PCA) of the normalized molecular features found under each treatment applied (n = 6). **(B)** Euler diagram of the common and uniquely molecular features with a differential and significant accumulation in each treatment (*P*_*adj*_ < 0.05). **(C)** Bubble diagram representing the up- and down-regulated molecular features found among the 568 molecular features uniquely and significantly changing under the combination of salinity + heat. Each bubble (i.e. molecular feature) is positioned in the chromatogram by its mass-to-charge (y-axis) and retention time (x-axis) and the size and darkness of one bubble represented the log_2_ and p-value, respectively of this molecular feature. The raw data of Figure 3C can be found in the supplemental material (Supplementary Table S2 and S3). **(D)** KEGG pathway enrichment analysis performed with the significantly up-regulated molecular features identified under the combination of salinity + heat. More details on these analyses can be found in the Materials and Methods section.

To identify important functions activated in response to each stress, an enrichment analysis of the significantly up-regulated genes was conducted using pathway annotations from the Kyoto Encyclopedia of Genes and Genomes (KEGG; **Supplemental Table S7**). A total of 27 pathways were found to be enriched (*P*_*adj*_ < 0.05) with these up-regulated genes across the three treatments (**Fig. 2C**). In line with the genes themselves, enriched pathways were largely unique, with all but three pathways (glutathione metabolism (sly00480), protein processing in the ER (sly04141), and spliceosome (sly03040)) being enriched in just one of the three treatments. Most interestingly, the salinity + heat treatment resulted in the upregulation of genes belonging to two main metabolic pathways, ASC and aldarate metabolism (sly00053) and arginine and proline metabolism (sly00330), which were not enriched in either of the individual stress treatments, suggesting that the combination of stresses induced specific changes in plant metabolism that in turn led to variation in the physiological responses of the plants.

### The salinity and heat combination showed a unique metabolic profile with the enrichment of specific pathways

A metabolomics study was carried out to identify molecular features that were common or unique to the simple or combined stresses and to validate the RNAseq results. Our main interest mainly resided in those that were specifically accumulated under the combination of salinity and heat. A total of 3,338 molecular features showed significant (*P*_*adj*_ < 0.05 and a log_2_ < -1 or log_2_ > 1) changes across the three stress conditions. Similar to the RNAseq analyses, each stress condition showed a unique metabolic profile (**Fig. 3A; Supplemental Tables S3 and S4**). Only 208 molecular features were commonly altered by all the treatments, which represented 6.30% of the total. When the combination of salinity and heat was applied, a total of 568 molecular features (17.19% of the total) were significantly and specifically accumulated with respect to the control (**Fig. 3B**). Salinity + heat caused reprogramming of multiple metabolic pathways, observed as a similar number of molecular features that were up- or down-regulated, 337 and 208, respectively, when compared to the control (**Fig. 3C**). Pathway enrichment analysis of the up-regulated molecular features revealed that four biochemical pathways (i.e., ASC and aldarate metabolism, purine metabolism, arginine and proline metabolism and arginine biosynthesis) were significantly altered in tomato under the combination of salinity and heat (**Fig. 3D and Supplementary Table S4, Supplementary Figs. S2 and S3)**. In agreement with the RNAseq data, the ASC and aldarate metabolism and the arginine and proline metabolism were among the most significantly enriched pathways.

### The integration of transcriptomics and metabolomics revealed that the proline and ASC pathways are interconnected for ROS homeostasis

The RNAseq and metabolomics data was combined with measurements of enzymatic activity to obtain a detailed picture of the changes in the ASC and aldarate, and arginine and proline metabolic pathways caused by the combination of salinity and heat stresses (**Fig. 4**). The first observation was that proline appears to be degraded in favor of 4-hydroxyproline and L-glutamate-5-semialdehyde accumulation, with the concomitant up-regulation of prolyl 4-hydroxylase (P4HA) and pyrroline-5-carboxylate reductase (PROC), as well as the down-regulation of proline dehydrogenase (PRODH). The accumulation of L-glutamate-5-semialdehyde was also likely derived from ornithine through the up-regulation of arginase (ARG) and ornithine aminotransferase (ROCD). In summary, proline was not differentially accumulated under the combination of salinity + heat as compared to controls. Instead, 4-hydroxyproline and L-glutamate-5-semialdehyde, two proline-derivative compounds, significantly accumulated in tomato leaves under stress combination.

**Figure 4.**
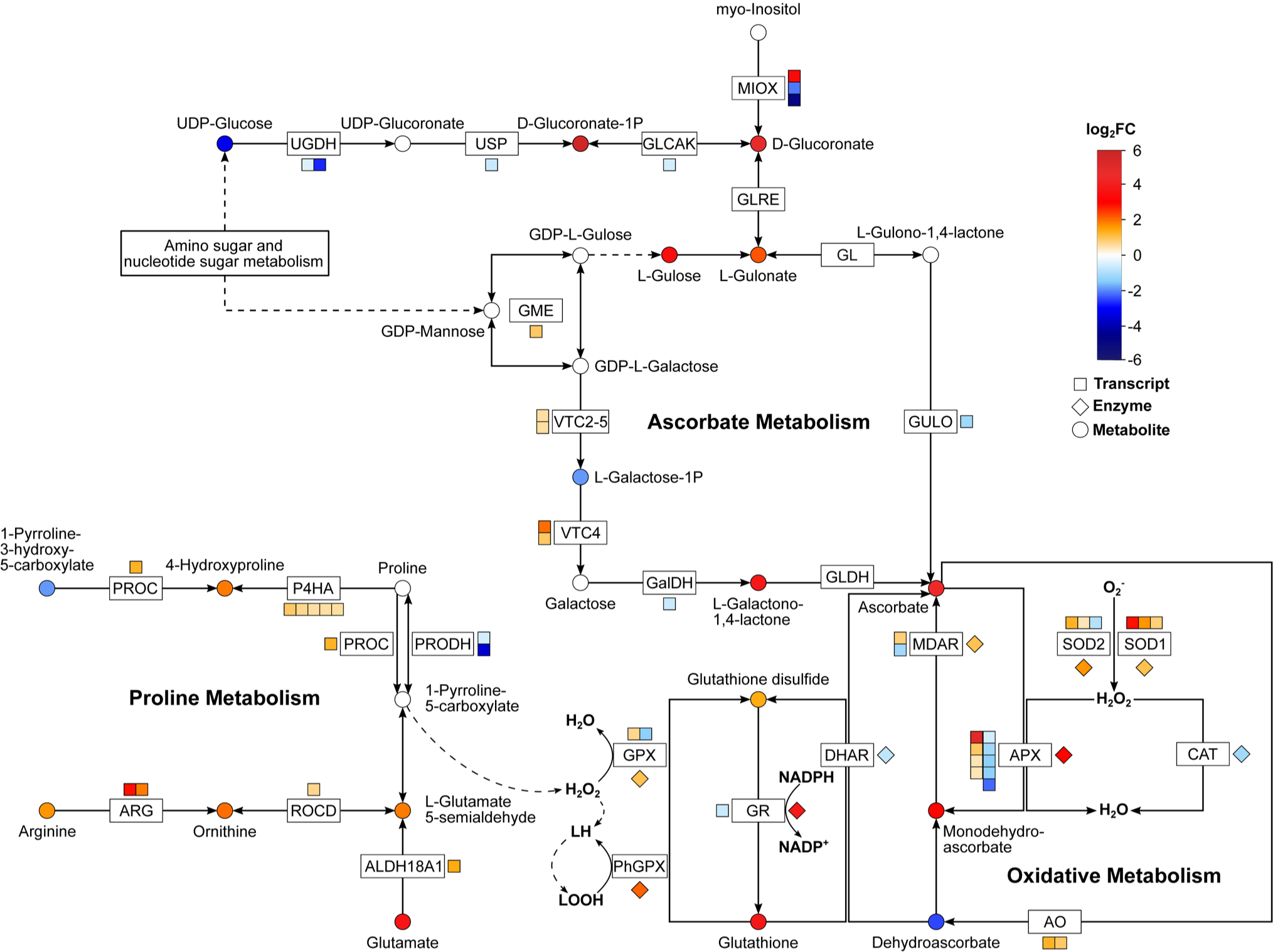
Schematic diagram of the metabolic interconnection between ascorbate, proline and oxidative metabolism in tomato plants. Log_2_(fold change) of the metabolite concentration (○), gene expression (□ □) or enzymatic activity (◊) obtained in tomato plants grown under the combination of salinity + heat after RNAseq, metabolomics or biochemical analyses were represented.

ASC significantly accumulated under combined salinity and heat stress in tomato. Its synthesis from UDP-glucose or myo-inositol results in the precursor D-glucuronate, which also increased under salinity + heat, in part due to the down-regulation of glucuronokinase transcript (GLCAK) through glucoronate-1P synthesis and to the up-regulation of one copy of the inositol oxygenase (MIOX). The levels of L-gulose and L-gulonate also increased under the combination of stresses, which seemed to favor ASC accumulation. The high ASC levels observed in tomato plants under stress combination could also be due to the degradation of the GDP-L-galactose and L-galactose-1P, since GDP-L-galactose phosphorylase (VTC2-5) and L-galactose 1-phosphate phosphatase (VTC4) were up-regulated under these conditions (**Fig. 4**). ASC is known to detoxify ROS through the Halliwell-Asada cycle. Remarkably, this pathway was highly represented among the differentially expressed genes and the significant molecular features altered by salinity + heat (**Figs. 2-3**). These results were also supported by the enzymatic activities of the proteins encoded by those transcripts (**Fig. 4**). Superoxide dismutases (SOD1 and SOD2), involved in cell ROS detoxification, were up-regulated at the transcript and activity levels, leading to the conversion of O_2_^-·^ to H_2_O_2_. Then, H_2_O_2_ can be detoxified by catalase (CAT) or by ASC peroxidase (APX) through the ASC/GSH pathway. CAT was not differentially expressed in the RNAseq analysis and the enzyme activity was inhibited by stress combination. Several APX homologs were up-regulated at the transcript level and, more importantly, its enzymatic activity was very high (log_2_ = 1.97, **Supplemental Table S5**) under salinity + heat. The APX activity generates monodehydroascorbate, which accumulated significantly in our experiments. Monodehydroascorbate spontaneously forms dehydroascorbate, which is reduced to ASC, once again through the action of dehydroascorbate reductase (DHAR), using glutathione (GSH) as a reducing agent. Tomato plants showed a significant accumulation of glutathione and monodehydroascorbate under combined salinity and heat stress, with a non-significant dehydroascorbate accumulation or DHAR activity. However, the MDAR enzyme, responsible for the regeneration of ASC, was up-regulated at the transcript and enzymatic levels. Lastly, the glutathione peroxidases GPX and PhGPX, responsible for the recovery of lipid peroxidation, were also up-regulated under stress combination. Our results are indicative of a connection between ASC synthesis and oxidative stress-proline metabolism, with the intersection between these pathways found at the 1-pyrroline-5-carboxylate level (**Fig. 4**).

### Plants subjected to salinity and heat combination showed lower oxidative damage than those under salinity alone

The proposed coordination between the proline, ASC, and redox pathways may improve the ability of the tomato plants to deal with ROS detoxification. Markers of oxidative stress were evaluated to determine if tomato plants subjected to stress combination displayed a more efficient antioxidant system than those plants grown under individual stresses (**Fig. 5**). Tomato plants under salinity had the highest levels of H_2_O_2_, with a significant 4-fold increase compared to control plants. However, when salinity and heat were applied in combination, it was found that the H_2_O_2_ content was about 50% lower than in the salinity treatment (**Fig. 5A**). A similar trend was observed for lipid peroxidation, an indicator of oxidative damage to cell membranes, with a maximum value found for salinity stress, and an intermediate value found for the salinity + heat stress combination (**Fig. 5B**). Thus, the stress combination appears to reduce oxidative damage with respect to the salinity treatment. In neither case, the differences between heat stress and control were statistically significant. Interestingly, the antioxidant capacity (**Fig. 5C**) obtained for plants subjected to salinity was the lowest among all treatments, with a reduction of up to 90% as compared to the controls. When salinity and heat were combined, the antioxidant capacity index was significantly lower than the control but 6-fold higher than salinity. Again, the heat treatment did not show significant differences with respect to the control. Protein oxidation values obtained for the four stress conditions were directly related to H_2_O_2_ and lipid peroxidation, with a positive and significant correlation (H_2_O_2_-protein oxidation: r = 0.992, *P*_*adj*_ < 0.001; lipid peroxidation-protein oxidation: r = 0.996, *P*_*adj*_ < 0.001). In short, our results indicated that ROS levels were lower when salinity and heat were applied jointly as compared to the salinity treatment alone, which was directly observed as a lower damage to the membrane lipids and to the cellular proteins under abiotic stress combination.

**Figure 5.**
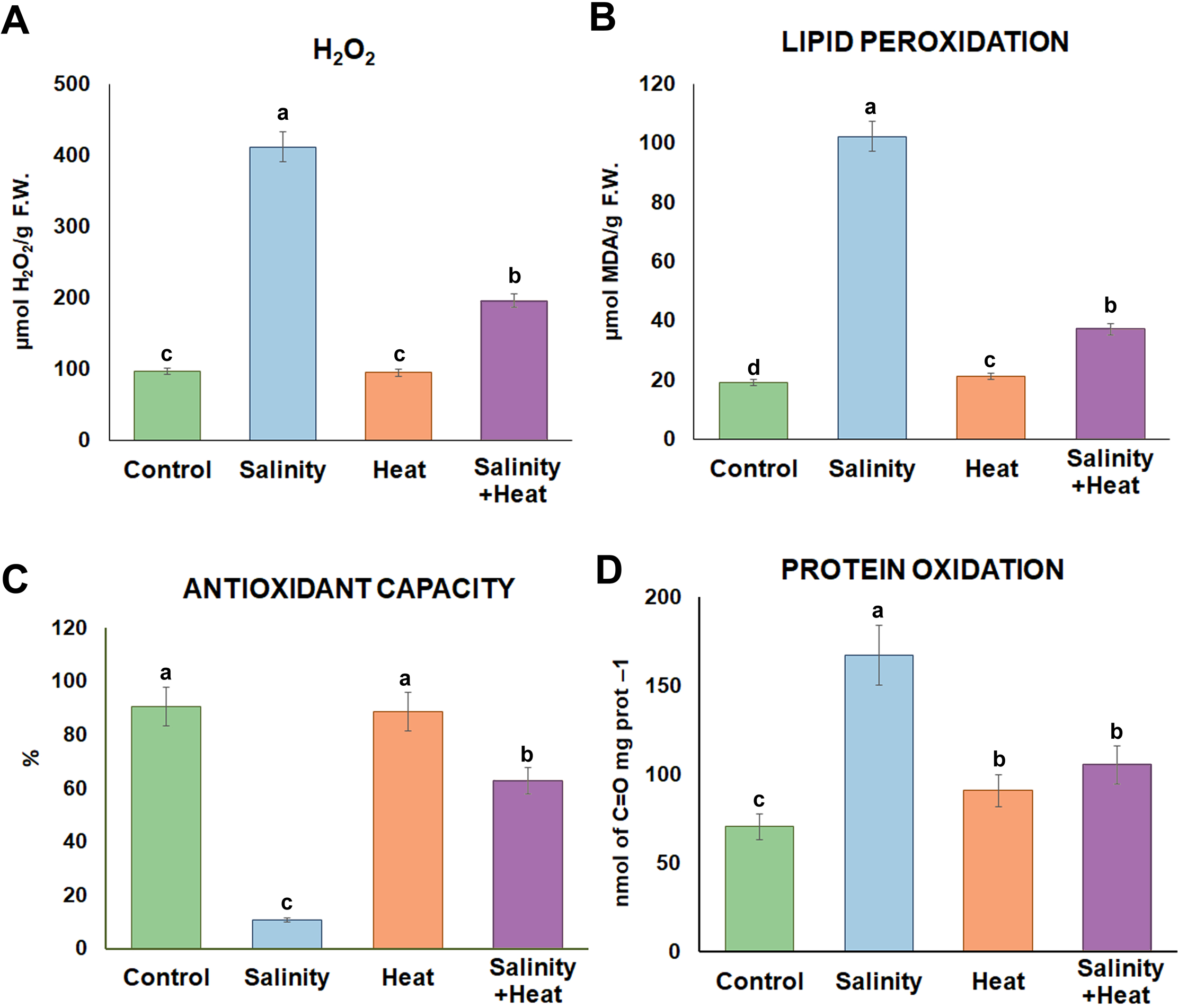
Oxidative metabolism-related markers measured in tomato leaves grown for 14 days under control, simple (salinity +heat) or combined (salinity+heat) stress. Values represent means ± SE (n = 9). Bars with different letters within each panel are significantly different at p < 0.05 according to Tukey’s test.

### The combined salinity and heat responses are associated with the upregulation of unique transcription factors families

The high specificity of the tomato plant responses to salinity + heat suggests that a tight regulatory control must be in place to rapidly and efficiently cope with the oxidative damage caused by these conditions. TFs are known to be key players in modulating the expression of genes involved in abiotic stress responses. TFs that may regulate the transcriptional responses to salinity, heat, and/or salinity + heat were identified by evaluating the promoter regions (1000 bp upstream from the transcription start site) of up-regulated genes from each stress condition for overrepresented cis-element motifs (**Fig. 6A**). Binding sites from a total of 46 TFs belonging to multiple gene families were found to be enriched (*P*_*adj*_ < 0.05) across all treatments. Of these, only 9 TFs were associated with all stress conditions (salinity, heat, and salinity + heat), including three Homeobox-Homeodomain-Leucine Zipper Protein (HB-HD-ZIP) TFs identified. The salinity + heat treatment exhibited five unique TFs, including three from the stress-related Zinc Finger Cysteine-2/Histidine-2 (C2H2) family.

**Figure 6.**
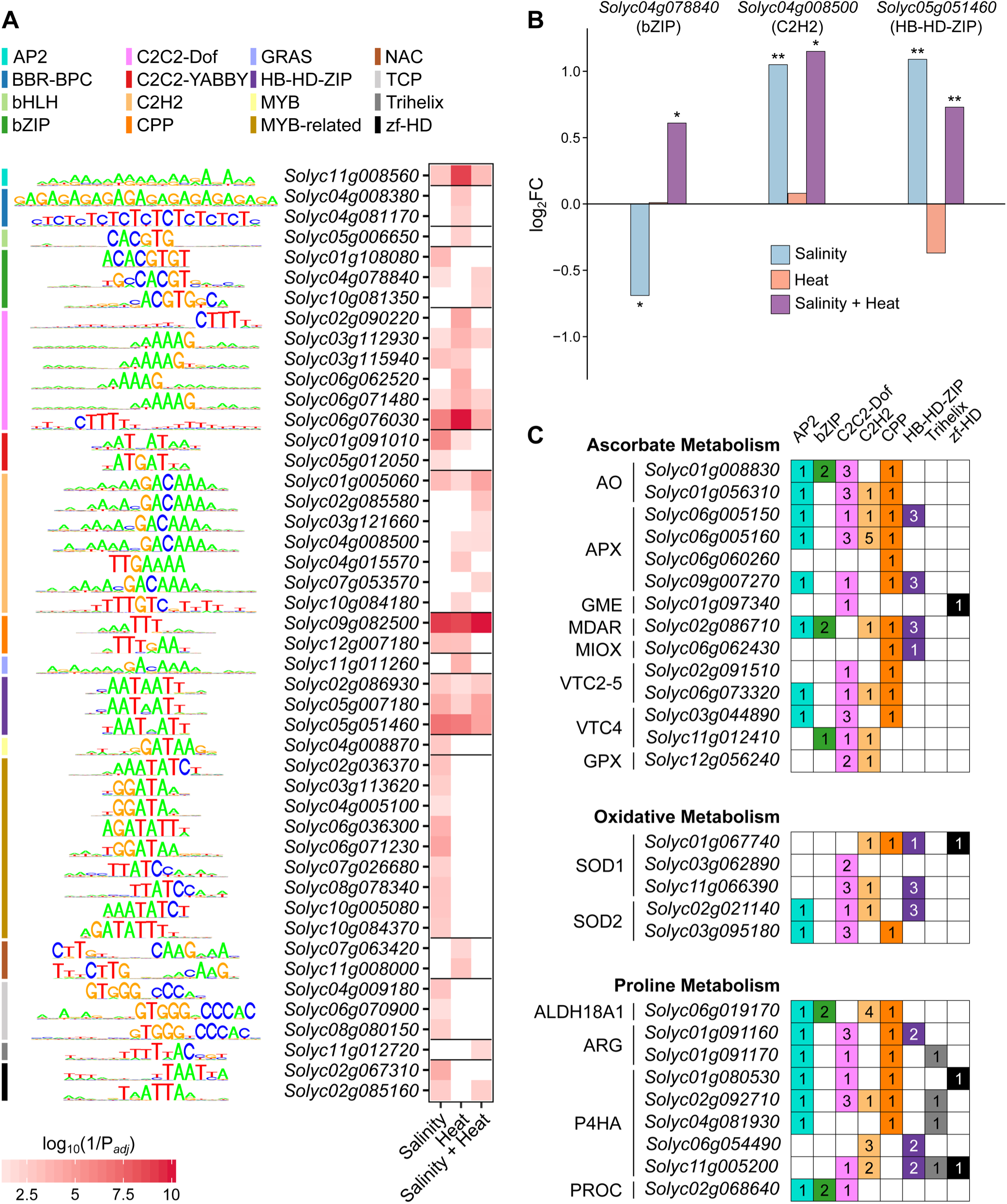
Cis-element enrichment results for up-regulated genes from each stress treatment.**(A)** Enrichment p-values for binding motifs corresponding to 46 TFs in each of the stress treatments. TFs are grouped and color coded by family, and a consensus diagram for the binding motif and gene accession is given for each. **(B)** Log_2_(fold change) of expression of three selected TFs in each stress treatment. **(C)** Counts of TF families overrepresented in genes up-regulated in the salinity + heat treatment from the ascorbate metabolism, oxidative metabolism, and proline metabolism families. Accessions and common abbreviations are given for each gene. Numbers in boxes refer to the count of TFs in that family with a match to that gene based on Analysis of Motif Enrichment results.

In contrast to the diverse enrichment results, most enriched TFs did not exhibit significant up-regulation themselves under their associated stress condition. For salinity + heat, only three TFs had significant (*P*_*adj*_ < 0.05) expression levels when compared to the controls (**Fig. 6B**). These TFs, one each from the basic Leucine Zipper Domain (bZIP), C2H2, and HB-HD-ZIP families, were also all differentially expressed in the salinity treatment, although the bZIP TF (*Solyc04g078840*) was down-regulated under this treatment, and up-regulated exclusively for salinity + heat. None of these three were differentially expressed under the heat stress alone. The sequences of the differentially expressed genes from the proline, ASC, and redox pathways (**Fig. 4**) were evaluated to determine which of the overrepresented cis-element motifs associated with salinity + heat were present in their promoters (**Fig. 6A**). Most of these genes include binding sites for TFs from the Apetala 2 (AP2), Dof zinc finger protein (C2C2-Dof), and Cysteine-rich Polycomb-like Protein (CPP) families, among others. Binding sites for the single CPP TF, which were highly enriched across all three stress conditions, matched to the promoters of genes from the proline, ASC, and oxidative metabolism pathways, including all four up-regulated copies of *APX* genes. Remarkably, binding sites for the single-enriched Trihelix TF, *Solyc11g012720*, were found only in the promoters of proline metabolism genes. Ultimately, these results suggest that specific sets of TFs coordinate the modulation of proline, ASC, and redox metabolism under salinity + heat stress, which should be further studied to validate their direct or indirect regulatory roles.

## DISCUSSION

In the present study, we demonstrated that the combination of heat stress with moderate salinity in tomato plants induced a specific physiological, biochemical, and molecular response that could not be deduced from a single stress application. From the physiological stand point, tomato plants under the combination of salinity and heat grew better than when salinity was applied as a sole stress, showing a significant increase in plant biomass. At the same line, plants under stress combination showed better photosynthetic performance and lower cellular oxidation than those growing under salinity, with the balance between these two processes necessary for both plant growth and adaptation to abiotic stress (Considine and Foyer, 2013; Woehle *et al*., 2017). Under salinity stress, ROS accumulation (measured as H_2_O_2_) likely induced damage to membranes and an increase in protein oxidation, which translated into a lower cell antioxidant capacity. These oxidative stress-associated processes may have caused the strong inhibition of photosynthesis and reduction of growth observed in plants subjected to the salinity treatment. Interestingly, when salinity was combined with heat, ROS were accumulated to a lesser extent, with the damage to membranes and proteins being also lower and maintaining an antioxidant capacity of over 60%, which was observed as plants with better growth rates than in the salinity conditions alone. These observations indicate that ROS could be produced in a lower quantity under stress combination than under salinity, and that ROS is being produced at the same level than under salinity, although their detoxification may be more efficient and/or effective under stress combination. Our results mainly support the latter possibility, in which the combination of salinity and heat induced the reprogramming of some important stress-related pathways, such as proline and ASC metabolism, facilitating their interconnectivity for a more efficient cellular ROS detoxification through the activation of oxidative metabolism.

Tomato plant responses to salinity and heat combination involved complex transcriptional networks and changes in metabolic fluxes. However, the modulation of the proline and ASC pathways was shown to be a strong and unique response to this stress combination. Interestingly, these metabolic pathways were not found to be significantly induced under salinity or heat when applied individually. Proline can protect cells from damage by acting as an osmoprotectant but also as a ROS scavenger (Szabados and Savouré, 2010; Narayanan and Govindarajan, 2012). Although proline metabolism was induced under the combination of salinity and heat, proline levels did not increase under these conditions, and instead, the derivatives 4-hydroxyproline and L-glutamate-5-semialdehyde were significantly accumulated. Several studies have pointed out that during stress recovery, proline is oxidized to provide the cell with a large amount of energy (one molecule of proline captures 30 ATP equivalents) (Verbruggen and Hermans, 2008; Zhang and Becker, 2015). Jaspers and Kangasjärvi (2010) showed that when salinity levels were increased, proline was used as a source of energy by plants, providing ATP and NADPH through its catalysis by the enzyme PRODH. This oxidation process increased the formation of ROS, activating the response signaling cascade generated by the oxidative stress (Jaspers and Kangasjärvi, 2010), and thus relating proline with the stress response mechanisms found in plants.

Our results pointed out to an interconnection between proline catalysis, ROS generation (due to stress conditions and proline degradation) and an upregulation of the oxidative metabolism. ASC metabolism was also up-regulated, as ASC is a necessary substrate to maintain oxidative metabolism active. It can be suggested that proline accumulation occurs early during the acclimation to stress combination and that its oxidation is a sign of stress recovery in these plants. However, we have previously reported that proline does not preferentially accumulate during the first 72 hours after tomato plants were subjected to the combination of salinity and heat stress (Rivero *et al*., 2014), which contradicts this idea. Instead, glycine-betaine was the osmolyte that was preferentially accumulated in tomato under these conditions. Thus, in this study, we propose and provide evidence that proline oxidation may be interconnected with glutathione redox homeostasis for efficient ROS scavenging. Recent publications have demonstrated that proline catabolism to P5C is induced in animal cells during cell infection (Tang and Pang, 2016). These authors proposed that PRODH and PROC act together to raise P5C levels and thus, govern ROS homeostasis. This mechanism is largely unknown in plants, although a similar hypothesis was proposed in *Arabidopsis thaliana* as a response to pathogen attack (Qamar *et al*., 2015). In a previous study by our research group (Rivero *et al*., 2014) we have also shown that PRODH and PROC were differentially up-regulated at the gene and protein levels under the combination of salinity and heat, whereas under salinity or heat applied individually these enzymes were down-regulated, thereby favoring proline accumulation. The significant enrichment of proline metabolism found in the analysis of the transcriptomics and metabolomics data and the potential role of proline intermediaries in ROS homeostasis, such as P5C, provide a strong argument for the role of proline oxidation in ROS signaling mechanisms under stress combination; however, we recognize that more research is needed to confirm this hypothesis.

ASC is one of the main compounds involved in plant oxidative metabolism through the Halliwell-Asada cycle, and genes and compounds found in this pathway were significantly induced under salinity and heat combination. Activities of the oxidative metabolism-related enzymes were determined to confirm the upregulation of the oxidative metabolism at the protein level, as well as to find whether or not this pathway was specifically regulated under the combination of salinity and heat, as previously shown for proline and ASC metabolism. Our research group, as well as other authors, have reported on the high activation of oxidative metabolism-related enzymes through a specific upregulation under the combination of salinity in tomato plants (Rivero *et al*., 2014; Martinez *et al*., 2018; García-Martí *et al*., 2019). The enzymatic activities assayed, together with the gene expression and the metabolites identified in our study, indicate the efficient detoxification of H_2_O_2_ through the Halliwell-Asada cycle, and a very active lipid recovery from oxidation thanks to PhGPX activity. These observations could explain that under salinity and heat combination, the oxidative markers measured (H_2_O_2_, lipid peroxidation, protein oxidation, and antioxidant capacity) in tomato plants were lower than under salinity as the sole stress.

Zandalinas *et al*. (2020) recently described that different combinations of abiotic stresses applied to *A. thaliana* plants resulted in unique transcriptional profiles and that their regulation by different TF families was also characteristic of each stress combination. In this report, the bHLH, MYB and bZIP TFs families were significantly induced under the combination of salinity and heat. Our results showed that some genes belonging to the bZIP TF family were differentially and uniquely regulated under the combination of salinity and heat in tomato plants (e.g., *Solyc10g081350*). Other TFs belonging to other stress-related families, such as C2H2 (e.g., *Solyc02g085580, Solyc03g121660, Solyc07g053570*) and Trihelix (e.g., *Solyc11g012720*), also showed this particularity under our experimental conditions. Most of these TFs families have been reported to be involved in the control of plant development, cell division, different physiological process, but also in abiotic responses of plants (Kaplan-Levy *et al*., 2014; Agarwal *et al*., 2019). For example, Agarwal *et al*. (2019) reported that the bZIP family was involved in the mitigation of several abiotic stresses (e.g., salinity, drought, heat or oxidative stress) and the increase in plant productivity under adverse conditions. The Trihelix TF family has been shown to be involved in the response to salinity and pathogen-related stresses, and in the development of trichomes, stomata, and the seed abscission layer (Kaplan-Levy *et al*., 2014). Numerous members of the C2H2-type zinc finger family have been shown to play a significant role in the plant’s response to different abiotic stresses and in plant hormonal transduction signals (Kiełbowicz-Matuk, 2012). Most of the information found in the literature regarding the C2H2 family has been for Arabidopsis, and very little is known about other plant species, including tomato. Hu *et al*. (2019) found that this family regulates many genes in response to some abiotic stress, and especially in response to heat stress in tomato plants. Most of the C2H2 genes that were up-regulated under heat stress in the report by Hu *et al*. (2019) were also differentially expressed in our transcriptomic analysis when heat was applied as the sole stress. However, the C2H2 identified in our study that was specifically up-regulated under salinity + heat was not listed in the study by Hu *et al*. (2019), again demonstrating the importance of studying stresses in combination. The TFs identified as being up-regulated under the combination of salinity and heat aligned with the promoter regions of many genes studied in this report, including those belonging to the proline, ASC, and oxidative metabolisms.

In summary, we showed that proline, ASC and oxidative metabolism are interconnected, with a tight coordination to maintain not only an optimal cellular redox balance, but also to trigger the proper signaling mechanisms responsible for inducing the plant’s acclimation to the combination of salinity and heat. In this process, proline oxidation is suggested to be used as the energy source needed for triggering the stress response, with subsequent ROS formation. At this point, oxidative metabolism enters the stage, with the upregulation of its main enzymes to maintain ROS at basal levels. One of the main limiting factors for maintaining the activity of the redox metabolic pathways is ASC abundance, which suggests the presence of a connection between ASC biosynthesis with oxidative metabolism and, most likely, with proline oxidation. Cellular basal levels of ROS could trigger downstream signaling mechanisms through the activation of particular TFs families, such as the trihelix, C2H2 and bZIP families, which in turn, may regulate the expression of genes involved in the reprogramming of different metabolic pathways, including those involved in proline, ASC, and redox metabolism (i.e., positive feedback loops). Future validation of the role of specific TFs families in the successful acclimation of plants to heat + salinity is necessary for developing breeding strategies for more resilient crops against abiotic stresses.

## Supplementary data

**Figure S1**. Total ion chromatogram extracted from UPLC-QTOF performed in 6 biological replications of tomato leaves subjected to control, salinity heat or the combination of salinity + heat.

**Figure S2**. Box-and-whisker plots of the compounds belong to the Ascobate, Alderate and oxidative metabolism with significant differences between salinity + heat treatment respect to control.

**Figure S3**. Box-and-whisker plots of the compounds belong to the Proline metabolism with significant differences between salinity + heat treatment respect to control.

**Table S1**. Raw, parsed and mapped reads of mRNA of all samples.

**Table S2** - **Sheet 1-**Comparison of salinity treated tomato plants against control plants. **Sheet 2-** Comparison of heat treated tomato plants against control plants. **Sheet 3-**Comparison of the salinity combined with heat treatment against control plants.

**Table S3**. Comparison of the peaks of each independent analysis with the aim of find common and specific peaks among all the treatments.

**Table S4**. Identified compounds in the Control vs Salinity+Heat peaks comparison related to the enriched pathway analysis results.

**Table S5**. Activities of the oxidative metabolism-related enzymes.

**Table S6**. Differential expression output from DESeq2 (Love et al., 2014).

**Table S7**. Enrichment of KEGG pathways in upregulated genes for each treatment.

## Data availability

The raw sequencing reads and the read mapping count matrices are available in the National Center for Biotechnology Information Gene Expression Omnibus database under the accession GSE152620. All data supporting the findings of this study are available within the paper and within its supplementary materials published online.

## Acknowledgements

This work was supported by the Ministry of Economy and Competitiveness from Spain (Grant No. PGC2018-09573-B-100) to R.M.R (Murcia, Spain), by the Ministry of Science, Innovation and Universities of Spain (Grant No. FPU16/05265) to M.L-D. (Murcia, Spain) and by start-up funds from the College of Agricultural and Environmental Sciences and the Department of Plant Sciences (UC Davis) to B.B-U (Davis, CA, United States). We sincerely acknowledge Mario G. Fon for proof-reading the manuscript. We also thank the Metabolomics Core of CEBAS-CSIC for the assistance with the analysis. All authors declare no commercial, industrial links or affiliations.

## Author contribution

RMR conceived, designed and supervised the experiments; ML-D, CJS and BB-U performed the RNAseq analysis. VM and RMR performed the metabolomic analysis. ML-D, TCM and RMR performed the biochemical analysis. ML-D and RMR performed the photosynthetic measurements. ML-D, BB-U and RMR wrote the paper. The authors declare that they have no competing interests.

